# RNA-Binding moonlighting function of metabolic enzymes reveals deep evolutionary roots in Cyanobacteria

**DOI:** 10.1101/2025.09.28.679013

**Authors:** S. Anudarsh, Dinesh Balasaheb Jadhav, Sougata Roy

## Abstract

RNA-binding proteins (RBPs) have emerged as key regulators of diverse physiological and metabolic processes in cells. Notably, many metabolic enzymes exhibit moonlighting RNA-binding functions, and a substantial fraction localize to chloroplasts, the primary hub of photosynthesis and cellular metabolic homeostasis. Since chloroplasts originated from free-living cyanobacteria, understanding the RBP repertoire in these ancient phototrophs holds particular evolutionary and functional significance. A comprehensive characterization of the cyanobacterial RBPome is still lacking. Here, we employed *Synechococcus elongatus* PCC 7942, a model cyanobacterium, to define its RBPome using an RNA-interactome capture approach. We identified 136 RBPs, of which nearly 30% are associated with metabolic pathways, a proportion notably higher than that observed in bacteria, algae, plants, flies, worms, or animals. Strikingly, several enzymes from core metabolic pathways, including glycolysis/gluconeogenesis, the TCA cycle, and the pentose phosphate pathway, that are known RNA binders in humans are also conserved as RBPs in cyanobacteria. We identified a wide array of proteins from the photosynthetic apparatus exhibiting RNA-binding activity, many of which are conserved across the green lineage. In silico structural alignments of RNA-binding metabolic enzymes with their NAD(P)-binding pockets, a potential site for RNA-binding, suggests a broad conservation of RNA-binding capacity of core metabolic enzymes across species. Recent discoveries have revealed that RNA-binding can modulate enzymatic activity. In this context, our findings suggest that RNA-mediated control of core cellular metabolic processes may be widespread in cyanobacteria and riboregulation might be an evolutionarily ancient mechanism, potentially tracing its origins back to cyanobacteria.

## Introduction

RBPs are highly conserved across taxa and represent one of the most functionally versatile classes of proteins, with roles spanning a wide range of physiological and metabolic processes (Lorković, 2009; Gerstberger et al., 2014; Liao et al., 2025). Traditionally, RBPs have been studied in the context of post-transcriptional regulation, where they associate with RNA molecules to influence their fate and function. Through specific interactions with RNA, RBPs govern essential processes such as splicing, polyadenylation, stability, localization, translation, and degradation (Glisovic et al., 2008). Collectively, these activities allow RBPs to act as central regulators of gene expression, fine-tuning the flow of genetic information and ensuring that cellular programs are precisely adapted to developmental, environmental, and metabolic demands.

Recent discoveries have uncovered an additional regulatory layer in which RNAs, rather than being merely passive targets, actively modulate the functions of RBPs (Hentze et al., 2018). This reciprocity becomes particularly intriguing with the realization that a large number of metabolic enzymes, across diverse biological systems, also function as non-canonical RBPs (Castello et al., 2015). In such cases, RNA binding is not limited to post-transcriptional regulation but can directly influence the catalytic properties of these enzymes. By modulating enzymatic activity, RNA– enzyme interactions can rewire cellular metabolic flux, thereby introducing a novel mechanism through which RNA exerts control over core metabolic and physiological processes. There is now growing recognition that RNA-binding proteins (RBPs) exhibit widespread moonlighting functions across biology, influencing processes from cellular energetics and core metabolic pathways to cell division, viral infection, and stem cell differentiation (Garcia-Moreno et al., 2019; Huppertz et al., 2022; Spizzichino et al., 2024; Rajagopal et al., 2025). Their pervasive roles in physiology and metabolism suggest that riboregulation is not a recent innovation but an evolutionarily ancient mechanism, fundamental to the temporal and functional organization of life. Keeping this in mind, in this study we employed the model cyanobacterium *S. elongatus* PCC 7942 and applied an unbiased phase separation–based RNA-interactome capture approach to define its RBP repertoire. The ancient evolutionary origins of cyanobacteria (Schirrmeister et al., 2015) provide a unique window into the primordial foundations of riboregulation. Notably, independent studies have already underscored the importance of RBPs in this phototrophic bacterium; for instance, Rbp2 has been shown to be essential for proper circadian clock function in *S. elongatus*, a premier model for circadian biology (McKnight et al., 2023). RBPs have been shown to be indispensable for cell survival under diverse stress conditions, including chill-light (Tan et al., 2011), cold (Mori et al., 1998), and salinity (Zhang et al., 2022). Beyond stress adaptation, several RBPs play essential roles in fundamental physiological processes such as photosynthesis (Hemm et al., 2025), thylakoid biogenesis (Wang et al., 2024), and heterocyst differentiation (Brenes-Álvarez et al., 2025), underscoring their central importance in both environmental resilience and core cellular functions in cyanobacteria. To date, very little is known about non-canonical RBPs in cyanobacteria, which could provide crucial insights into the evolutionary origins and consequences of their moonlighting function. Emerging evidence suggests that certain metabolic enzymes can “moonlight,” switching between their canonical catalytic functions and RNA-binding activities. This functional duality appears to be regulated, at least in part, by temporal post-translational modifications, which may determine when an enzyme engages in catalysis versus RNA interaction (Huppertz et al., 2022; Neusius et al., 2022; Spizzichino et al., 2024). Such regulation provides a potential mechanism for dynamically integrating metabolic flux with RNA-based control across the circadian cycle. In our lab, we found that RNA binding to metabolic enzymes indeed exhibits subjective day–night dynamics (Jadhav and Roy, 2025a). Notably, many of the enzymes displaying such daily temporal variation in RNA association are localized to the chloroplast (Jadhav and Roy, 2025a), a relic of the free-living cyanobacteria.

In our investigation, we identified 136 RBPs in *S. elongatus* PCC 7942, representing nearly 5% of the total proteome, consistent with RNA interactome capture studies across species. Remarkably, about 30% of these RBPs are associated with metabolic pathways, the highest proportion reported so far in any species using this approach. The RBP repertoire of *S. elongatus* includes enzymes from core primary metabolic processes such as glycolysis/gluconeogenesis, the TCA cycle, the pentose phosphate pathway, and sucrose and starch metabolism. Notably, the RNA-binding functions of several key enzymes show striking evolutionary conservation, extending from cyanobacteria to humans. Similarly, components of the photosynthetic apparatus extensively exhibited moonlighting RNA-binding functions. Several of these key photosynthetic proteins are conserved across the green lineage, and our *in silico* analysis further suggests that this RNA-binding activity may also be preserved in red and brown algae. Together, our findings demonstrate that RNA-binding moonlighting functions are ancient, raising the strong possibility that riboregulation represents an evolutionarily conserved mechanism for fine-tuning cellular metabolic flux.

## Results

### RNA-interactome capture catalogues the RBPome of the cyanobacteria *S. elongatus*

To comprehensively catalogue RNA-binding proteins (RBPs) in the cyanobacterial model *S. elongatus* PCC 7942, we employed an orthogonal organic phase separation method, which enables the capture of RNA–protein complexes independent of poly(A) tail presence. Successful RBP enrichment in this approach critically depends on effective zero-distance crosslinking of RNA to its interacting proteins. UV crosslinking at 254 nm has been widely adopted for this purpose due to its specificity and efficiency (Urdaneta and Beckmann, 2020). Given the absence of prior global RBPome studies in cyanobacteria, we first optimized the UV dose required for effective crosslinking. *S. elongatus* cells were cultured under 12 h:12 h light: dark cycles in a Percival algal growth chamber (AL41L4) at 30±1 °C for at least 7 days. Cells were harvested at an OD_750_ and samples were collected across four distinct time points spanning the day–night cycle (Figure 1A), to capture RNA–protein interactions that may have a temporal regulation. For each time point, biological replicates were divided into crosslinked (CL) and non-crosslinked (NC) controls (n=4 per condition).

**Figure 1:**
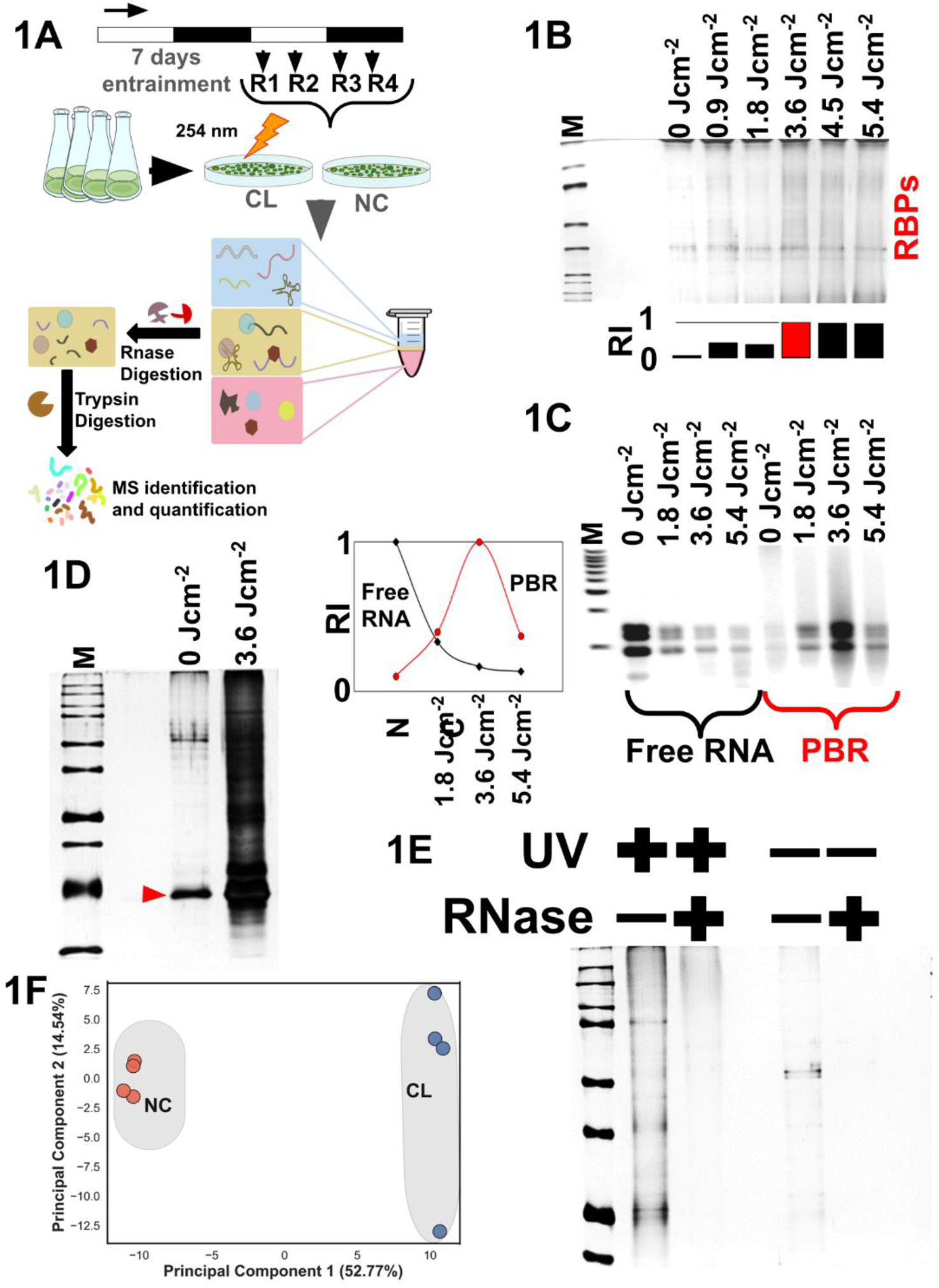
RNA-interactome capture in *S. elongatus* 7942 and its RBPome. (1A) The overall schematics of our capture approach and the downstream processing. (1B) Silver-stained gels containing the captured RBPs were loaded, with different lanes representing varying doses of crosslinking. The relative intensity (RI) of the protein bands in each lane was quantified using ImageJ software. Multiple horizontal cross-sectional panels were analyzed in ImageJ, and the final intensity value was obtained by summing the individual intensities across all panels. (1C) PBRs and free RNAs from crosslinked and non-crosslinked cells were resolved on a horizontal RNA gel containing ethidium bromide. Relative intensity (RI) was quantified using ImageJ, with rRNA band intensity serving as the reference. (2D) Histogram showing the proportion of RBPs associated with metabolic pathways as percentage. RBPs were identified either from RBPbase or from individual published datasets. The count of RBPs involved in specific metabolic pathways was assigned using KEGG’s inbuilt mapper tool, and their distribution was represented as proportions across pathways. (2E) RNase treated and untreated extracts from cross-linked and non-crosslinked samples were loaded onto a 10% PAGE supplemented with 10% SDS and visualized by silver staining. (1F) The PCA plot using the peptide intensity obtained from cross linked and non-crosslinked samples after mass spectrometry analysis.

To determine the optimal UV dose for crosslinking, cells were exposed to varying doses (0, 0.9, 1.8, 3.6, 4.5, and 5.4 J cm⁻²), followed by RBP extraction and silver staining of equal protein volumes on 10% SDS–PAGE (Figure 1B). The dose yielding the highest protein recovery, 3.6 J cm⁻², was selected for subsequent experiments (Figure 1D). This was independently validated by quantifying free RNA and protein-bound RNA (PBR) fractions (Figure 1C); the optimal dose should yield maximal PBR with minimal free RNA, which has been previously referred as crosslinking efficiency (Queiroz et al., 2019; Villanueva et al., 2020). Both analyses converged on 3.6 J cm⁻² as the optimal crosslinking dose. To validate the specificity of the method, we included RNase digestion controls. If the proteins enriched in CL samples were genuine RBPs, RNase treatment prior to extraction would abolish their recovery. Indeed, silver staining of RNase-digested samples showed almost no detectable protein bands, confirming the specificity of RNA-dependent enrichment (Figure 1E).

Following optimization, RBPs were extracted from all CL and NC biological replicates. In parallel, additional pooled samples were processed under identical conditions to generate an empirical spectral library for subsequent mass spectrometry analysis (Figure 1A). All protein samples were digested with trypsin and subjected to quantitative SWATH-MS (Sequential Window Acquisition of All Theoretical Mass Spectra) at the IGIB mass spectrometry facility. For spectral library generation, a standard 1% false discovery rate (FDR) was applied at the spectrum, peptide, and protein levels, resulting in the identification of 3399 spectra, 1923 peptides, and 353 unique proteins. Using this spectral library, SWATH-MS quantified protein intensities across all CL and NC samples. Comparative analysis revealed pronounced differences in total protein intensities between CL and NC replicates, consistent with selective enrichment of RBPs in the crosslinked condition.

### Metabolic enzymes with RNA-binding function in cyanobacteria

a. *S. elongatus* possesses a robust circadian clock that regulates a wide array of cellular processes (Cohen and Golden, 2015). Based on this, we hypothesized that RNA–RBP interactions in this organism would also exhibit distinct temporal partitioning. To ensure we do not miss RBPs that show temporal binding patterns, we collected samples at four different time points across the day and night, using them as independent biological replicates in our analysis. To identify cyanobacterial RNA-binding proteins (cyRBPs), we applied a stringent threshold of >2-fold enrichment in crosslinked (CL) versus non-crosslinked (NC) samples, with a −log10(p-value) > 1. Proteins showing fold changes between 1.5 and 2 were considered enriched “candidate RBPs”. In total, from the four independent replicates, we identified 136 cyRBPs. This recovery falls within the typical range reported for RBP identification in other systems. For instance, in mammals, RBPs typically constitute about 5% of the total proteome, as estimated from RNA-interacting proteins in the UniProt reference set. In comparison, recovery rates from plants, flies, worms, and *E. coli* (Stenum et al., 2023) via RIC are often lower than 5%, although an orthogonal phase separation method in *E. coli* (Queiroz et al., 2019) has achieved up to 8% RBP recovery (Figure 2B). STRING analysis of the 136 cyRBPs revealed five major functional clusters, with translation emerging as the largest, comprising 73 proteins( Interestingly, carbon fixation, photosynthesis and antioxidant activity)(Supplementary Figure 2). Along with this, the functional classification validated the effectiveness of the RIC approach, showing strong enrichment for canonical RBPs linked to Gene Ontology (GO) terms such as ribosome, RNA and rRNA binding, RNA metabolism, translation, and tRNA aminoacylation(Figure 2C). Notably, components of the photosynthetic machinery were also enriched, along with GO categories related to primary metabolic processes, gene expression, and antioxidant activity.

**Figure 2:**
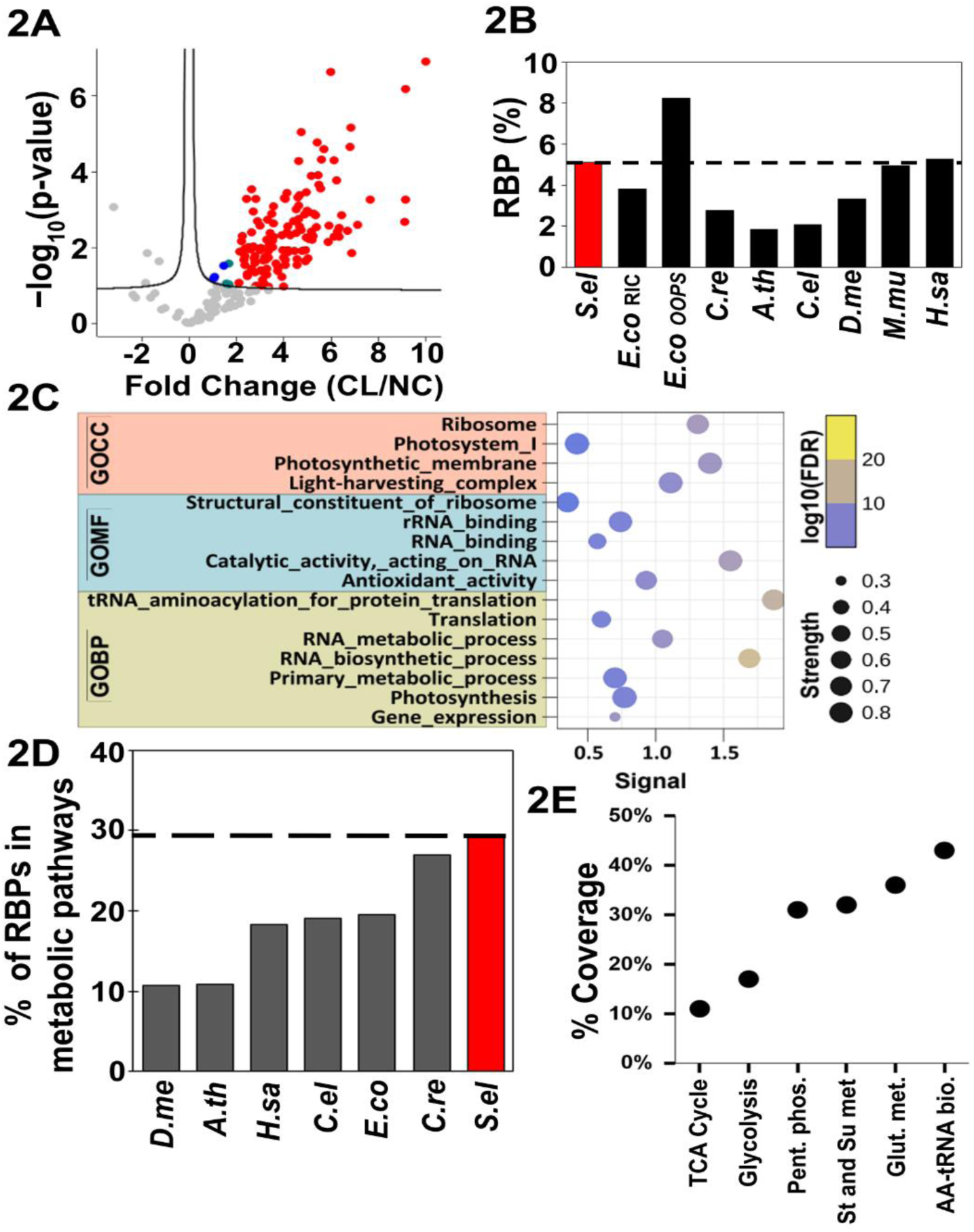
The moonlighting function of RNA-binding is rampant in metabolic enzymes of *S. elongatus*. (2A) Volcano plot showing identified cyRBPs and candidate cyRBPs. Proteins with FDR < 0.05 and log2(FC CL/No-CL) ≥ 2 are classified as cyRBPs (red), while proteins with 1 ≤ log2(FC CL/No-CL) < 2 and FDR < 0.05 are designated as candidate cyRBPs (blue). (2B) Proportion of RBPs expressed as a percentage of total proteins across organisms. RBP numbers were obtained either from published studies or retrieved from RBPbase (https://apps.embl.de/rbpbase/), while total protein counts were derived from the reference proteomes available in UniProt. The dotted line indicates the reference percentage of RBPs for *S. elongatus*. (2C) Gene Ontology (GO) enrichment analysis for the cyanobacterial RNA-binding proteins (cyRBPs) was performed using the STRING database (https://string-db.org/). The analysis covered the three major GO categories: Cellular Component, Biological Process, and Molecular Function. STRING’s built-in enrichment algorithm was applied to identify significantly overrepresented terms relative to the reference proteome, and enrichment values (including FDR-adjusted *p*-values) were obtained directly from the platform. (2D) Representative silver-stained gel showing the final capture of RBPs prior to mass spectrometry. The red arrow indicates the RNase band, the enzyme used to digest RNAs before RBP loading. (2E) Dot plot showing the proportion of S. elongatus enzymes from primary metabolic pathways exhibiting RNA-binding moonlighting activity. The values are expressed as coverage, calculated as the ratio of RNA-binding enzymes to the total number of enzymes in each pathway. Both numerator and denominator values were obtained from the KEGG pathway database.

To evaluate the involvement of cyRBPs in metabolic pathways, we used KEGG Mapper by inputting corresponding gene IDs. This enabled pathway-level assignment of the identified RBPs. For cross-species comparison, we conducted similar analyses using RBP datasets from other organisms, obtained from RBPbase (https://apps.embl.de/rbpbase/) and published studies. These were also mapped to metabolic pathways via KEGG Mapper. Strikingly, in *S. elongatus*, about 30% (40 of the 136) of the identified RBPs were associated with metabolic pathways, the highest percentage observed among all species analyzed(Figure 2D). This prompted a deeper investigation into the primary metabolic pathways enriched for RNA-binding enzymes. We calculated the proportion of RNA-binding enzymes per pathway, expressed as a percentage relative to the total number of enzymes in each respective pathway(Figure 2E). Our findings indicate that cyRBPs are represented across all major cellular metabolic pathways, a trend that appears to be conserved across diverse species.

### Conserved RNA-Binding functions of core metabolic enzymes from Cyanobacteria to Humans

We present a detailed investigation into the evolutionary conservation of RNA-binding activity across species, from cyanobacteria to humans, with a particular focus on key metabolic enzymes and the pathways they regulate. In *S. elongatus*, four enzymes from the glycolysis/gluconeogenesis pathway were identified as RNA-binding proteins (RBPs)(Figure 3A). Notably, one of them, fructose bisphosphatase (FBPase), plays a dual role: it is a critical enzyme in gluconeogenesis and a rate-limiting step in the Calvin cycle (Miyagawa et al., 2001). FBPase activity plays a pivotal role in enhancing the regenerative capacity of the Calvin cycle by facilitating through the pentose phosphate pathway (PPP) the production of ribulose-1,5-bisphosphate (RuBP), the primary CO₂-acceptor molecule. This, in turn, is essential for sustaining carbon fixation and directly contributes to increased photosynthetic efficiency (Miyagawa et al., 2001). The discovery of RNA-binding activity in FBPase introduces the possibility of riboregulation as a regulatory layer influencing both glucose metabolism and photosynthetic efficiency in cyanobacteria. Furthermore, we found in a previous study in our laboratory that the RNA-binding function of FBPase is conserved in *Chlamydomonas reinhardtii* (Jadhav and Roy, 2025a), suggesting that this regulatory mechanism may be evolutionarily conserved across green algae. Among the other three enzymes, RNA-binding activity of fructose bisphosphate aldolase (FBA) and phosphoglycerate kinase (PGK) appears to be conserved from cyanobacteria to humans. Both enzymes are central players in core carbon metabolic pathways, catalyzing key steps in glycolysis, gluconeogenesis, and the Calvin-Benson cycle in photosynthetic organisms. The functional significance of their RNA-binding activity remains unclear: it is yet to be determined whether this feature modulates RNA metabolism, or conversely, if RNA binding serves to temporally fine-tune the enzymatic activity of FBA and PGK. Similar to SHMT1, which exhibits a dual role in both translation regulation and being regulated by RNA, these enzymes may also engage in such moonlighting functions (Spizzichino et al., 2024), suggesting an intricate layer of riboregulation embedded within metabolic control. The fourth enzyme is the NAD(P)-dependent glyceraldehyde-3-phosphate dehydrogenase, commonly referred to as GAPDH2 in cyanobacteria. In vegetative cells of cyanobacteria, GAPDH2 is the predominant isoform, while the expression of the NAD-dependent GAPDH1, typically the most widespread GAPDH isoforms across species is barely detectable (Valverde et al., 2001). GAPDH1’s RNA-binding function has been extensively documented across diverse systems, where it plays crucial roles in regulating mRNA stability, translation, and mediating cellular responses to stress and inflammation (Galván-Peña et al., 2019; Garcin, 2019; Shamloo et al., 2024). Intriguingly, RNA-binding appears to compete with the cofactor (NAD⁺) binding in GAPDH (Nagy and Rigby, 1995), suggesting a potential regulatory switch that toggles the enzyme between its metabolic and RNA-associated roles. Furthermore, post-translational modifications (PTMs) such as oxidation (Arutyunova et al., 2003), acetylation (Huppertz et al., 2022; Neusius et al., 2022), malonylation (Galván-Peña et al., 2019) or glycation (Sofronova et al., 2021) have been shown to modulate this balance, enhancing or inhibiting RNA-binding affinity, thereby adding another layer of control over its dual functionality. Given that GAPDH2 can bind to NAD+ or NADP+ with equal efficiency it is logical to state that GAPDH2 in *S. elongatus* 7942 will follow similar RNA-binding functions and undergo analogous regulatory mechanism upon post translational modification.

**Figure 3:**
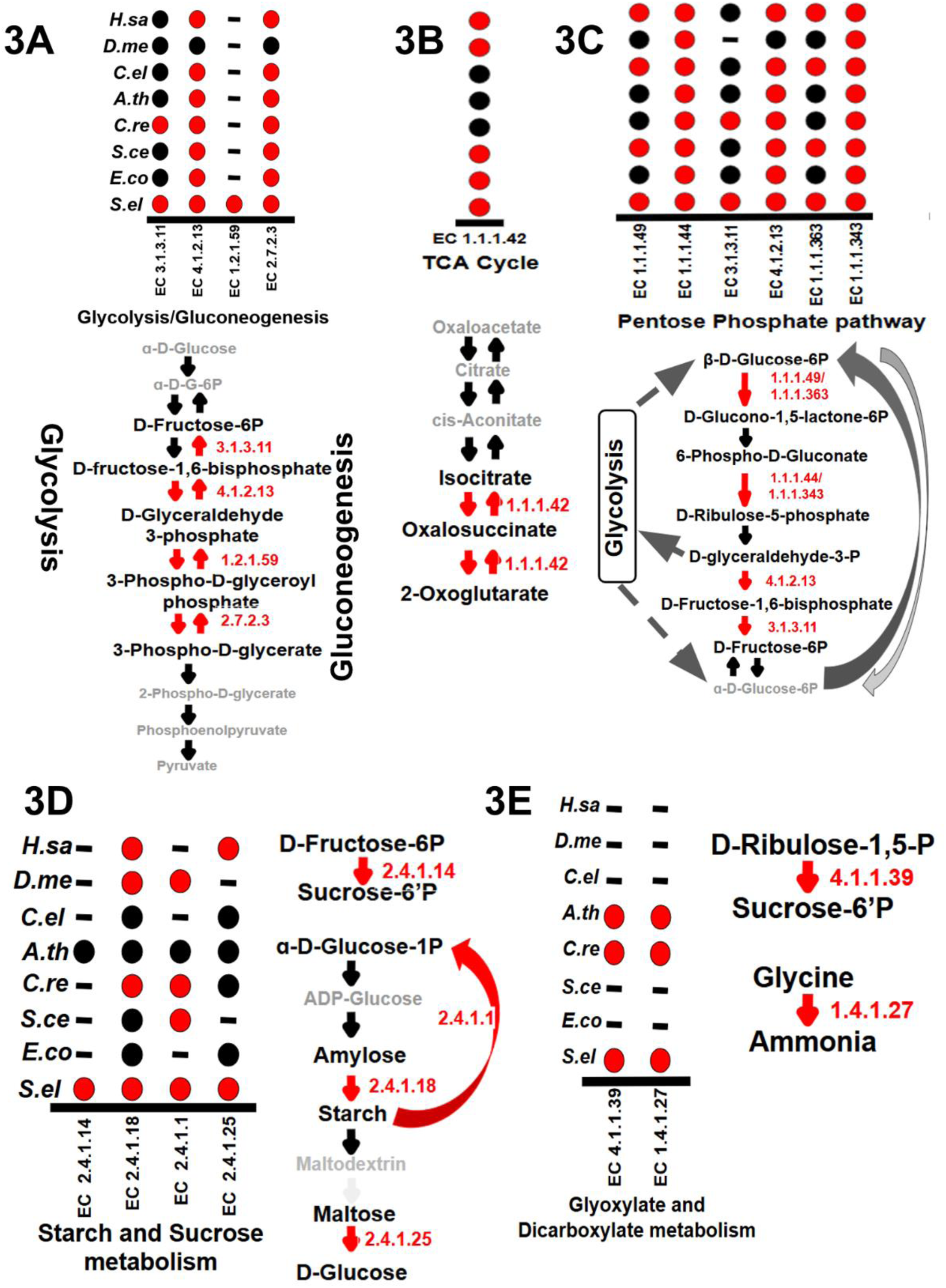
RNA-binding moonlighting function of metabolic enzymes remains conserved from cyanobacteria to humans. Red dots and arrows indicate RNA-binding metabolic enzymes. Black dots and arrows represent metabolic enzymes that do not exhibit RNA-binding activity. Black dashes denote the absence of the corresponding enzyme in that species. Top panel: RNA-binding metabolic enzymes from the primary metabolic pathways in *S. elongatus* and their conservation status across taxa. Bottom or side panel: Specific reactions within the pathway catalyzed by the RNA-binding metabolic enzyme(s). (3A) Glycolysis/gluconeogenesis, (3B) Tricarboxylic acid cycle (citric acid cycle), (3C) Pentose phosphate pathway (PPP), (3D) Starch and sucrose metabolism, and (3E) Glyoxylate and dicarboxylate metabolism.

2-oxoglutarate serves as a central carbon skeleton for amino acid biosynthesis, particularly for glutamate and glutamine, thereby playing a critical role in linking carbon and nitrogen metabolism (Huergo and Dixon, 2015). In both plants and bacteria, this metabolite is essential for efficient ammonium assimilation and the subsequent synthesis of nitrogenous compounds (Lancien et al., 2000; Doucette et al., 2011). 2-oxoglutarate is a cofactor for many dioxygenases involved in DNA and histone demethylation, impacting gene expression and cell fate (Fletcher and Coleman, 2020; Frost et al., 2021). The enzymatic interconversion of isocitrate to 2-oxoglutarate, catalyzed by NADP-dependent isocitrate dehydrogenase, is a critical step in the tricarboxylic acid (TCA) cycle. This reaction is pivotal not only for carbon flux through central metabolism but also for the generation of NADH/NADPH, which directly fuels cellular energetics by supporting ATP synthesis. Thus, this step represents a vital nexus between metabolic activity and energy homeostasis. Notably, we identified the NADP-dependent isocitrate dehydrogenase, responsible for this reaction, as an RNA-binding protein conserved across species. This observation suggests a potential layer of riboregulation could govern central metabolic flux, which remains conserved in cyanobacteria and humans.

The pentose phosphate pathway (PPP) is an essential anabolic route that operates parallel to glycolysis, primarily responsible for generating pentose sugars for nucleotide biosynthesis and reducing equivalents in the form of NADPH, which is vital for reductive biosynthetic processes and cellular redox homeostasis (Stincone et al., 2015; Rashida and Laxman, 2021). In the oxidative branch of PPP, glucose-6-phosphate is oxidized to ribose 5-phosphate through a series of enzymatic steps. In addition to the RNA-binding glycolytic enzymes described above that also participate in the PPP, we identified two key oxidative PPP enzymes, glucose-6-phosphate dehydrogenase (G6PD) and 6-phosphogluconate dehydrogenase (6-PGD), as RNA-binding proteins(Figure 3C). These enzymes catalyze the sequential reactions converting glucose-6-phosphate to ribulose 5-phosphate, underscoring a potential regulatory layer where RNA-binding may fine-tune PPP flux, linking metabolic and post-transcriptional regulatory networks. D-ribulose-5-phosphate (Ru5P) is a key metabolic intermediate whose levels are critical for efficient carbon fixation, energy production, and biosynthetic processes. As a central metabolite in the pentose phosphate pathway and Calvin-Benson cycle, the regulation of Ru5P is essential for maintaining metabolic balance in both photosynthetic and non-photosynthetic organisms. In plants and microbes, tight control over Ru5P ensures an optimal supply of ribose sugars for nucleotide synthesis and regeneration of ribulose-1,5-bisphosphate (RuBP) for CO₂ fixation. Perturbations in Ru5P homeostasis can therefore profoundly affect cellular growth, photosynthetic efficiency, and the biosynthesis of essential macromolecules.

Sucrose functions as an osmoprotectant against salt stress in *S. elongatus* (Desplats et al., 2005; Lin et al., 2020). Beyond its physiological role, sucrose is also a valuable commercial product and a key starting material for bioethanol production. We identified the enzyme sucrose-phosphate synthase (SPS), which catalyzes the conversion of D-fructose 6-phosphate and UDP-glucose to sucrose-6-P, to be an RNA-binding protein(Figure 3D). Salt stress activates expression of the rate-limiting SPS, thereby enhancing sucrose production (Desplats et al., 2005; Lin et al., 2020). Given that RNA-binding has been shown to modulate the activity of various enzymes, this discovery suggests a potential regulatory mechanism through which RNA binding could fine tune sucrose flux. This also opens up intriguing possibilities for RNA switches to fine-tune sucrose metabolic output via riboregulation.

### Rampant RNA-binding activity of photosynthetic proteins and their emergence in cyanobacteria

In the green lineage, ferredoxin–NADP⁺ reductase (FNR) plays a crucial role as a terminal electron acceptor from ferredoxin (Fd) in the photosynthetic electron transport chain (Carrillo and Ceccarelli, 2003). This is used to catalyze the reduction of NADP⁺ into NADPH. The NADPH produced in this reaction serves as a key reducing power for various metabolic processes, most notably the Calvin–Benson cycle, where it drives the fixation of atmospheric CO₂ into organic compounds. We discovered that NADP-dependent ferredoxin reductase (FNR) exhibits RNA-binding activity in cyanobacteria, a feature conserved throughout the green lineage(Figure 4A). In eukaryotes, similar RNA-binding can occur through the NAD⁺-binding domains of NAD-dependent MDH2 and GAPDH (Nagy and Rigby, 1995; Noble et al., 2024). Such interactions can modulate the catalytic activity of these enzymes, making it particularly intriguing to investigate the functional implications of RNA-binding in FNR. Structure-based alignment of FNR across various phytoplankton species suggests that RNA-binding may also occur in brown and red algae(Figure 4B). The role of FNR as an RNA-binding protein (RBP) appears to be highly significant in photosynthetic organisms. *S. elongatus* offers a simple and tractable model to investigate this function in detail. In GAPDH, the NAD⁺-binding pocket has been identified as a potential RNA-binding region (Nagy and Rigby, 1995). Positively charged residues such as arginine and lysine are known to be critical for RNA–protein interactions (Pina et al., 2018; Hofweber and Dormann, 2019), and sequence alignment of NAD⁺ pockets across the aforementioned species reveals strong conservation of two such positively charged amino acids within the pocket(Figure 4D). Many proteins from Photosystem I (PSI), Photosystem II (PSII), and the photosynthetic electron transport (PET) chain in *S. elongatus* exhibit RNA-binding activity(Figure 4A). Many of these RNA-binding interactions are conserved in *C. reinhardtii*, *A. thaliana*, or both, underscoring their potential evolutionary significance. Notably, in an earlier study from our laboratory, we discovered that PsbD, a core component of PSII responsible for light-driven water splitting, binds RNA in a time-of-day–dependent manner, with distinct binding patterns during the subjective day and night (Jadhav and Roy, 2025b). This rhythmic RNA-binding suggests a possible layer of riboregulation in photosynthesis, wherein RNA-protein interactions modulate photosynthetic function over the circadian cycle.

**Figure 4:**
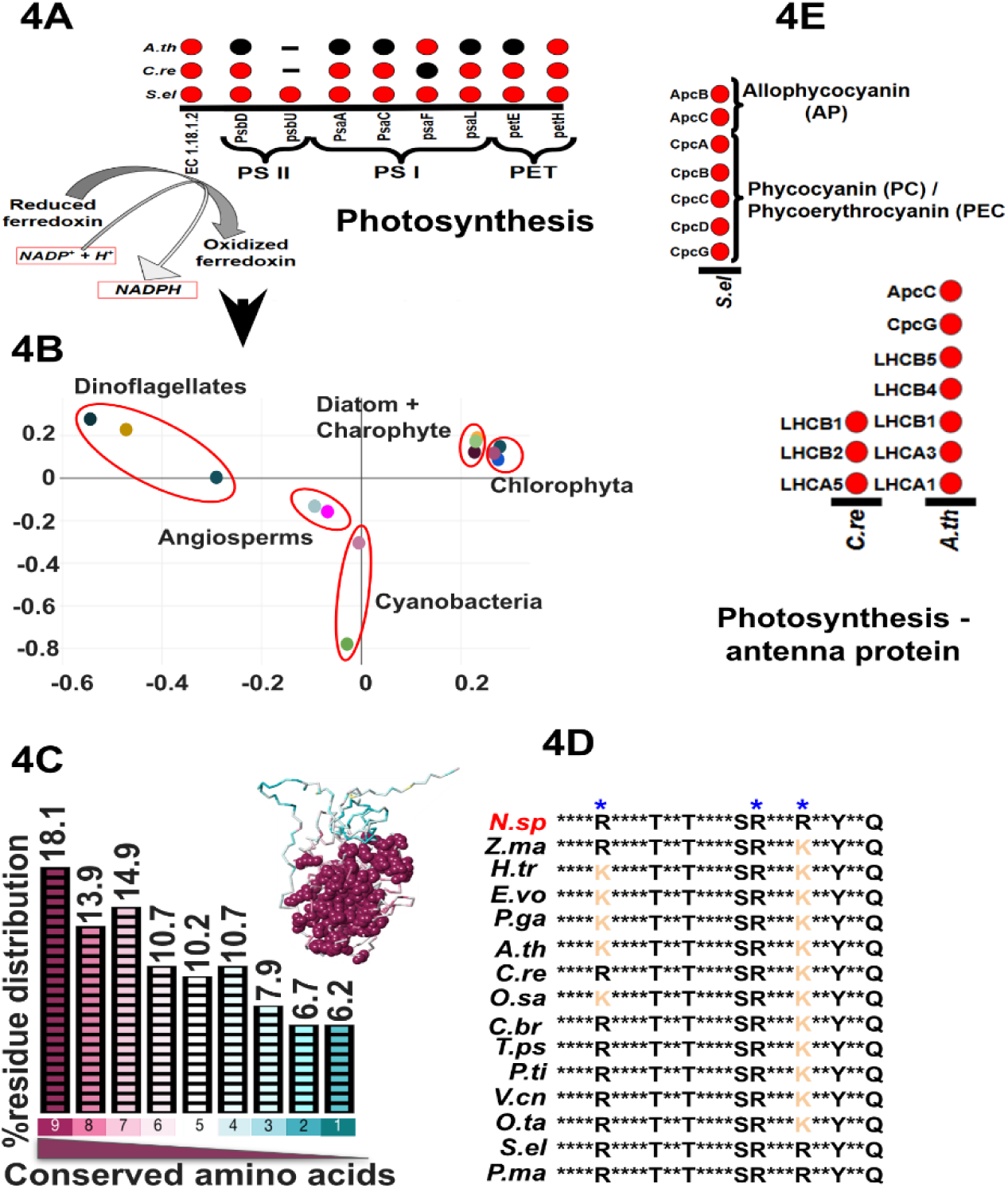
RNA-binding activity of photosynthetic apparatus proteins is conserved across the green lineage. Red dots indicate RNA-binding photosynthetic apparatus proteins. Black dots represent proteins that do not exhibit RNA-binding activity. Black dashes denote the absence of the corresponding protein in that species. **(4A)** RNA-binding activity across PSI, PSII, and the photosynthetic electron transport (PET) chain in *S. elongatus* and its conservation in chlorophytes and plants **(4B)** Correspondence analysis of structure-based alignment of ferredoxin–NADP (+) reductase (FNR; EC 1.18.1.2) across photosynthetic species. Each point represents a protein structure from the indicated species. Relative distances between points reflect structural similarity, derived from pairwise DALI Z-scores. The accession numbers and species names of representative FNR proteins used in this analysis are provided in Supplementary Table 2. **(4C)** This histogram shows the amino acid conservation profile of FNR from 41 different species using the ConSurf server (http://consurf.tau.ac.il/ ). The amino acid residue conservation percentage is calculated based on the input multiple sequence alignments (MSA) and phylogenetic relationships. The MSA was performed using MUSCLE alignment option from MEGA 11 (version 11.0.13) with default parameters. Position-specific conservation scores were computed based on the empirical Bayesian method, and the amino acid substitution model was chosen by default. Each bar of the histogram shows the percentage of residues falling into each conservation score, where the score of 9 with maroon represents the most conserved positions. The intermediately conserved positions are marked with white, having a score of 5 and the most variable positions are coloured in turquoise and have a score of 1. **(4 Graphical)** The conservation-based colours are projected on to the 3D structure of the protein. Only the most conserved amino acid scores are shown as balls and rest of the colour-coded sticks represent the following scores of conservations. **(4D)** The crystallised FNR structure of *Nostoc sp. PCC 7119* (pdb 1GJR) with its ligand Nicotinamide-adenine-dinucleotide phosphate was used as a reference and determined the interacting amino acids from PoseView and PoseEdit(https://proteins.plus/ ). Following which, those amino acids are represented as letters. The highlighted blue star represents the conserved cationic amino acids that are interacting with the ligand.

In cyanobacteria, the antenna of the photosynthetic light-harvesting complex consists of phycobilisomes (PBS), which are composed of phycobiliproteins (PBP) covalently bound to phycobilin chromophores. In *S. elongatus*, we identified several PBPs, including phycocyanins (PC), allophycocyanins (APC), and phycoerythrocyanins (PEC) that exhibit RNA-binding activity(Figure 4E). Similar RNA-binding functions have previously been reported in *Chlamydomonas reinhardtii* (Jadhav and Roy, 2025a) by our laboratory and in *Arabidopsis thaliana* (Reichel et al., 2016; Zhang et al., 2023) by others, suggesting that this RNA-binding function is conserved across diverse photosynthetic lineages.

## Discussion

Cyanobacteria, which emerged around 2.3 billion years ago, are among the most ecologically and evolutionarily significant microbes in Earth’s history, having played a central role in the planet’s oxygenation and the evolution of aerobic life. Although individual RBPs in cyanobacteria have been implicated in diverse and critical physiological processes, ranging from adaptation to environmental stresses such as salinity and temperature, to the regulation of the circadian clock, there has been no systematic effort to map the global RBPome of this important class of organisms. Such a comprehensive analysis is essential to uncover the full repertoire of RNA–protein interactions in cyanobacteria, which may reveal conserved regulatory strategies and novel post-transcriptional control mechanisms that shaped the evolution of photosynthetic life. Furthermore, the recent discovery of riboregulation, where RNA binding can modulate the catalytic activity of metabolic enzymes, has intensified our interest in exploring the extent of its conservation and its evolutionary origins. Understanding whether this regulatory mechanism is deeply rooted in cyanobacteria could provide key insights into how RNA–protein interactions have shaped both metabolism and gene expression since the earliest stages of evolution of aerobic life and photosynthesis.

The circadian clock in *S. elongatus* is known to influence every aspect of its biochemistry, physiology, and metabolism (Pattanayak and Rust, 2014; Cohen and Golden, 2015; de Barros Dantas et al., 2023). To capture potential temporally active RBPs, we collected cells at multiple time points across both the day and night. For downstream analyses, these time points were grouped into day and night biological replicates. By capturing cyanobacterial RNA-binding proteins (cyRBPs), we uncovered key insights into the prevalence of RNA-binding enzymatic functions and their potential evolutionary origins. These findings support the hypothesis that RNA-binding “moonlighting” functions are ancient and conserved across the tree of life, from cyanobacteria to humans, highlighting riboregulation as a fundamental and widespread regulatory mechanism.

The enrichment of pathways such as translation, RNA and rRNA binding, RNA metabolism, and ribosome in our dataset provides strong confidence in the quality and biological relevance of our RBP identification. Moreover, we detected RNP1, RNP2, and RNP3, well-characterized canonical RNA-binding proteins, within our repertoire, further validating the approach. Notably, the RNA-binding protein Rbp2, which we identified, was recently reported to be essential for the proper localization of the KaiC clock protein in *S. elongatus*(McKnight et al., 2023), underscoring the critical role of RBPs in the circadian clock function of this cyanobacterium. Our study not only delivers the first comprehensive RBPome of a cyanobacterium but also uncovers several intriguing candidates that open new avenues for exploring the interplay between RNA regulation, photosynthesis, metabolism, and circadian rhythms in prokaryotic phototrophs.

The discovery of RNA-binding as a moonlighting function in chloroplast proteins is well documented in plants and green algae, prompting us to investigate its potential ancestral origin in cyanobacteria. Our analyses in *S. elongatus* revealed that RNA-binding as a secondary (“moonlighting”) activity is not only widespread but also likely conserved across diverse cyanobacterial lineages. Chloroplast RBPs are distributed across the entire photosynthetic apparatus, encompassing components of the light-harvesting PBS, core proteins of PSI and PSII, as well as key factors within the PET chain.

Strikingly, among all organisms in which the RNA-binding proteome has been comprehensively characterized, *S. elongatus* displayed the highest proportion (almost 30%) of metabolic pathway proteins exhibiting RNA-binding activity. The RNA-binding activity of proteins spans across all major conserved primary metabolic pathways, including glycolysis/gluconeogenesis, the TCA cycle, the pentose phosphate pathway, sucrose and starch metabolism, and antioxidant defense. Antioxidant activity is among the earliest biochemical adaptations, likely arising in response to the surge in atmospheric molecular oxygen during the Great Oxidation Event (GOE). The proteins involved in this pathway binding RNA is particularly intriguing, suggesting a potential regulatory layer that warrants deeper investigation.

This suggests that RNA-protein interactions in metabolic enzymes are a deeply rooted feature of cyanobacterial physiology, potentially predating the endosymbiotic origin of chloroplasts. Such extensive coupling between metabolism and RNA regulation in *S. elongatus* highlights its value as a model system to uncover the evolutionary and mechanistic basis of riboregulation in photosynthetic organisms.

## Methods

### Culture conditions for *S. elongatus* PCC 7942

a. *S. elongatus* wild-type strain PCC 7942 (a kind gift from Prof. Susan Golden, UCSD) was inoculated from modified BG-11 agar plates into 500 mL of fresh BG-11 medium. Cultures were grown in an algal chamber (Percival AL41L4) under 12 h light/12 h dark (LD 12:12) cycles at 30±1 °C, with continuous shaking in a light intensity of 100 µmol m^-2^ s^-1^. Cells were entrained under these conditions for 16 days prior to sampling. Cell density was monitored spectrophotometrically at OD₇₅₀ (Clariostar, BMG Labtech). When cultures reached OD₇₅₀ ≈ 1, for each replicate, ∼1 × 10¹⁰ cells were harvested under diurnal conditions. In total, four replicates were collected (two from day and two from night), each derived from independent starter plates and considered as independent biological replicates. Cells were pelleted by centrifugation at 8000 X g for 10 min at 30 °C, washed once with fresh BG-11 medium, and immediately processed for downstream analyses

### RNA-Interactome capture

a. For each replicate, two samples of ∼1 × 10¹⁰ cells were collected: one subjected to crosslinking (CL) and the other processed as a non-crosslinked (No-CL) control.
b. **Crosslinking of cells:** For crosslinking, cell pellets were resuspended in 10 mL BG-11 and evenly spread onto 90 mm Petri plates. Plates were positioned 5 cm below the UV source, and irradiated at 254 nm using a UVP CL-1000 crosslinker (Analytik Jena). Different UV doses (0.9, 1.8, 3.6, 4.5, and 5.4 Jcm^-2^) were tested to determine the optimal conditions for RNA–protein crosslinking. Each UV-dose were applied as an increment of 0.45 Jcm-² with one-minute intervals between doses. The corresponding No-CL control cells were kept at room temperature during this procedure.
c. **Cell lysis:** After crosslinking, both CL and No-CL cells were transferred to Falcon tubes and centrifuged at 8000 g for 10 min at 4 °C. Pellets were resuspended in 200 µL lysis buffer [10 mM Tris-HCl (pH 7.4), 500 mM NaCl, 1 mM EDTA (pH 8), 5 mM DTT, 1 mM PMSF, and 40 U/µL RNase inhibitor(Invitrogen, Cat. No. 10777-019) in HPLC-grade water] in 1.5 mL microcentrifuge tubes preloaded with 300 µL of 0.1 mm pre-chilled zirconia beads(BioSpec Products Cat. No. 11079101z). Cells were disrupted at 70 Hz for three cycles of 30 s each, with 10 s cooling intervals, at 4 °C in a bead beater (Servicebio). Lysates were immediately processed or kept at −80 °C after adding Tri-Xtract (G-Biosciences, Cat. No.786-652).
d. **Isolation of RNA-binding proteins (RBPs) and Protein-Bound RNAs (PBRs) from the RNA-interactome:** This procedure was performed following a previously published protocol. Briefly, after centrifugation at 8000 g for 10 min at 4 °C, 200 µL of cell lysate was mixed with 2 mL Tri-Xtract and 400 µL chloroform, vortexed for 15 s, and centrifuged at 15,000 g for 10 min at 4 °C. Free RNA was isolated from the aqueous phase according to the Tri-Xtract manual. Protein-bound RNAs (PBRs) and RBPs were recovered from the interphase after two successive washes with 1 mL Tri-Xtract and 200 µL chloroform, with purification of the interphase after each wash. The RNA interactome was then precipitated by adding 1 mL of methanol, vortexing for 15 s, and centrifuging at 15,000 g for 10 min at 4 °C. This step was repeated once more. The resulting pellet was resuspended in 100 µL 100 mM TEAB (pH 8.5; Sigma-Aldrich, Cat. No. T7408) containing 1% SDS and heated at 95 °C for 7 min.

To isolate RBPs, samples were cooled and treated with 4 µg RNase A (Thermo Fisher Scientific, Cat. No. EN0531) and 10 U RNase T1 (Thermo Fisher Scientific, Cat. No. EN0541), followed by incubation for 16 h at 37 °C with shaking at 400 rpm (ThermoMixer, Thermo Fisher). After digestion, 1 mL Tri-Xtract and 200 µL chloroform were added, vortexed for 15 s, and centrifuged at 15,000 g for 10 min at 4 °C. The aqueous and interphase fractions were discarded, and 350 µL of the organic phase was transferred to a fresh tube. RBPs were precipitated by adding nine volumes of absolute ethanol and centrifuging at 15,000 g for 10 min at 4 °C. The pellet was washed with 1 mL of 70% ethanol, vortexed, and centrifuged under the same conditions. After discarding the supernatant and air-drying for 5 min, RBPs were resuspended in 40 µL 100 mM TEAB containing 0.1% SDS. Protein yield was ∼5–6 µg, as determined by Bradford assay. RBPs were separated using 10–12% SDS–polyacrylamide gels. For each lane, 8 µL of protein sample (8.63 ng/µL in TEAB with 0.1% SDS) was mixed with 1× Laemmli buffer, heated at 95 °C for 10 min, and loaded onto the gel. After electrophoresis, gels were fixed in 50 mL of fixing solution (50% ethanol, 5% acetic acid in double-distilled water) for 30 min on a rocker and stained with silver. Stained gels were visualized using a ChemiDoc imaging system (Bio-Rad).

For PBR isolation, the methanol-precipitated interphase was used. An enzyme-buffer mix containing 300 µL of Proteinase K buffer (10 mM Tris-HCl, pH 7.5; 1 mM EDTA, pH 8.0), and 12 U of Proteinase K (Merck, Cat. No. 1.24568.0100) was prepared and incubated at 50 °C for 15 min at 400 rpm. This mix was then added to the interphase and incubated for 4 h at 50 °C with 400 rpm to digest proteins. Following digestion, 300 µL of Tri-Xtract and 60 µL of chloroform were added, mixed vigorously for 15 s, and centrifuged. After phase separation, free RNA from the aqueous phase was recovered by adding 50ul of sodium acetate (3M) and 600ul of absolute isopropanol and incubated on ice for 15 minutes. It was centrifuged at 15,000g for 10 minutes at 4 °C. Supernatant was discarded and centrifuged again with 1ml of 70% ethanol. Pellets were air-dried for 5 min at room temperature and resuspended in DEPC-treated water. RNA concentration was measured using a NanoDrop spectrophotometer (Thermo Fisher Scientific), and RNA quality was assessed by agarose gel electrophoresis.

For the RNase digestion assay, we followed a protocol described in a previous study (Reference). Briefly, for CL, 10,000 million cells were irradiated with a 3.6 Jcm^-2^ UV dose, and NC samples served as controls. RNase T1 and A treatment for digestion was done for 16 hours at 37 ℃. The purified cyRBPs were resolved on 12% SDS-PAGE gels, and subsequent silver staining revealed distinct protein profiles in RNase-treated and untreated samples.

### SWATH-MS Proteomics

a. **Sample preparation for the Spectral library**

A spectral library is a prerequisite for efficient peptide identification and subsequent quantification while using SWATH-MS. For the spectral library generation, the same protocol of sample preparation and downstream processing was followed, but with a higher number of cells for each sample. 60,000 million entrained cells were collected from the same time points as described before. The cells were crosslinked as above, and after lysis, combined with 5 ml of Tri-Xtract and 1 ml of chloroform in a new 15 ml falcon tube. The interphase was retained and washed twice by repeating the procedure as mentioned earlier. Then 5 ml of methanol was added to the interphase to precipitate the RNA-interactome, and centrifuged twice as mentioned above. The pellets were air dried, and 500 µl of TEAB with 1% SDS was used to resuspend them. This was heated at 95 ℃ for 7 minutes. After cooling, 50 µl of 10X RNase buffer(100mM Tris-HCl (pH 7.5), 1.5 M NaCl,0.5%(vol/vol) NP-40 and 5 mM DTT) and 22 µg RNase A, and 50 µg RNase T1 were added and incubated at 37 ℃ for 16 hours at 400 rpm. After the incubation, 1 ml Tri-Xtract was added to the samples and followed the same steps as mentioned above.

**b.** Reduction, Alkylation and Trypsin digestion

The purified RBPs were reduced with 10 mM Dithiothreitol (DTT) at 56 ℃ for 45 minutes. Following this, 20 mM Iodoacetamide was added for the alkylation at room temperature and incubated in the dark for 45 minutes as per the previously described protocol elsewhere (). After that, a 16-hour enzymatic digestion with trypsin was performed at 37 degrees Celsius at 400 rpm with the enzyme-to-substrate ratio of 1:20 (trypsin: proteins), the enzymatic reaction was halted with 0.5%Trifluroacetic acid (TFA). Peptides were desalted with a C18 spin column (Pierce Cat. No. 89851) according to the manufacturer’s protocol. The peptide concentration was estimated using the Bradford assay and Nanodrop absorbance.

**c.** Spectral ion library generation

The trypsin-digested peptides were fractioned into 8 fractions by a cation exchange cartridge and with the aid of ascending concentration of ammonium formate buffer (35 mM-350mM ammonium formate, 30%v/v ACN, and 0.1% formic acid; pH=2.9). C-18 ZipTips were used for the cleaning up of the peptides from each fraction. Analysis of each fraction was performed on a quadrupole-TOF hybrid mass spectrometer (TripleTOF 6600, SCIEX) coupled to an Eksigent NanoLC-425 system. Optimized source parameters were used, curtain gas and nebulizer gas were maintained at 25 psi and 20 psi, respectively, the ion spray voltage was set to 5.5 kV, and the temperature was set to 250 degrees Celsius. About 4 μg of peptides were loaded on a trap column, and online desalting was performed with a flow rate of 10 μL per minute for 10 minutes. Peptides were separated on a reverse-phase C18 analytical column (ChromXP C18, 3 µm 120 Å, Eksigent, SCIEX) in a 57-minute-long buffer gradient with a flow rate of 5 µl/minute using water with 0.1% formic acid (buffer A) and acetonitrile with 0.1% formic acid. Analyst TF 1.7.1 Software was used for data acquisition. A 1.3-sec instrument cycle was repeated in high sensitivity mode throughout the entire gradient, consisting of a full scan of MS spectrum (400–1250 m/z) with an accumulated time of 0.25s, followed by 20 MS/MS experiments (100–1500 m/z) with 50 msec accumulation time each, on MS precursors with charge state 2+ to 5+ exceeding a 120-cps threshold. The rolling collision energy was used, and the former target ions were excluded for 15 seconds.

**d.** SWATH-MS data acquisition

The in-solution digested samples from the two cycles were analyzed in SWATH-MS mode on the same instrument with similar LC gradient and source parameters as DDA runs. A SWATH-MS method was created with 100 precursor isolation windows, defined based on precursor m/z frequencies in a DDA run using the SWATH Variable Window Calculator (SCIEX), with a minimum window of 5 m/z. Accumulation time was set to 250 msec for the MS scan (400–1250 m/z) and 25 msec for the MS/MS scans (100–1500 m/z). Rolling collision energies were applied for each window based on the m/z range of each SWATH and a charge 2+ ion, with a collision energy spread of 5. The total cycle time was 2.8 sec.

**e.** Data Analysis

A merged database search for DDA runs was performed using Proteinpilot^TM^ Software 5.0.1 (SCIEX) against the S. elongatus proteome from UniprotKB (UP000889800, with 2,657 protein entries). The Paragon algorithm was used to get protein group identities. The search parameters were set as follows: sample type-identification, cysteine alkylation-iodoacetamide, and digestion-trypsin. The biological modification was enabled in ID focus. The search effort was set to ‘Thorough ID’ and the detected protein threshold [Unused ProtScore (Conf)] was set to >0.05 (10.0%). False discovery rate (FDR) analysis was enabled. Only proteins identified with 1% global FDR were considered true identification. The group search result file from Proteinpilot^TM^ Software was used as a spectral ion library for SWATH analysis.

SWATH peak areas were extracted using SWATH 2.0 microapp in PeakView 2.2 software (SCIEX), and shared peptides were excluded. SWATH run files were added, and retention time calibration was performed using peptides from abundant proteins. The processing settings for peak extraction were a maximum of 10 peptides per protein, 5 transitions per peptide, >95% peptide confidence threshold, and 1% peptide FDR. The XIC extraction window was set to 5 min with a 50 ppm XIC Width. All information was exported in the form of MarkerView (.mrkvw) files. In MarkerView 1.2.1 (SCIEX), protein area data were normalized; the data normalization strategy used was total area sum normalization, and further analysis was performed using Python, R, MaxQuant Perseus, and Microsoft Excel.

### Proteomic data normalisation and statistical analysis

Missing values in the raw intensity data were imputed using the *MinDet* function implemented in Python (v3.11.7) with the pandas package (v2.2.2). Data normalization was performed using quantile normalization implemented in Normalyzer(). The CL intensities followed a normal distribution. The normalized intensity values from non-crosslinked (No-CL) and crosslinked (CL) samples were uploaded into MaxQuant Perseus (v2.0.11.0). To identify RNA-binding proteins (RBPs), a two-sided *t*-test was performed using quantile-normalized data from four biological replicates. The parameters were set to a false discovery rate (FDR) of 0.05, s0 = 0.1, with 250 randomizations. Proteins meeting the criteria of FDR < 0.05 and log2(FC CL/No-CL) ≥ 2 were classified as cyRBPs, while those with 1 ≤ log2(FC CL/No-CL) < 2 and FDR < 0.05 were designated as candidate cyRBPs. Downstream analyses, including principal component analysis (PCA), Pearson correlation, hierarchical clustering, and volcano plots, were performed using either Python or the corresponding functions within Perseus. To gain functional insights into the enriched RBPs, the dataset was uploaded to the STRING (https://string-db.org/ ) database for gene ontology (GO) enrichment analysis. The analysis revealed significantly overrepresented GO terms across three major categories: cellular component, biological process, and molecular function. This approach allowed us to systematically annotate the RBPs, providing a broader understanding of their potential cellular roles, regulatory pathways, and molecular interactions that were plotted using python.

All the enriched RBPs were queried on the STRING and the resulting protein networks were clustered by K-means clustering with the number of nodes as eight. The resultant network clusters were imported to Cytoscape (have to cite the original paper) for visualization.

### Percentage enrichment of RBP across species

The RBP descriptive information table of each species was downloaded from the EMBL RBPbase (https://apps.embl.de/rbpbase/). From the RBP descriptive information table the gene identifiers were selected only if the RBP is present in any of the species specific studies. Those species specific RBP’s corresponding gene names were collated and used for the KEGG mapper search (https://www.genome.jp/kegg/mapper/search.html). The KEGG mapper provided the quantitative categorization of metabolic and other pathways.

### Sequence and Structure analysis across species

For the structural alignment analysis of Ferredoxin NADP reductase (FNR). 13 photosynthetic species sequences were chosen from Uniprot and NCBI. Top search results with FNR of chloroplastic type were selected. The species and their Uniprot/NCBI accession IDs are provided in the supplementary table 2.

For 6-phosphogluconate dehydrogenase, decarboxylating (6-PGD) 8 sequences were used for the structural analysis. The sequences used are listed in supplementary table 2. Three of the structures were obtained from RCSB PDB.

FNR and the other 5 PGD structures were predicted using AlphaFold3 webserver (https://alphafoldserver.com/) with the default setting (Abramson et al., 2024). All the model 0 structures were chosen from the five predicted structures. The CIF files were converted to PDB files using a web based tool (https://project-gemmi.github.io/wasm/convert/cif2pdb.html). The structural comparisons were performed in the DALI web server (http://ekhidna2.biocenter.helsinki.fi/dali/) (Holm, 2022) using all against all structural comparisons.

The conservation of amino acids were evaluated based on the phylogenetic relationship between the 41 FNR and 28 6-PGD sequences using the Consurf webserver (https://consurf.tau.ac.il/consurf_index.php)(Ashkenazy et al., 2016). All the species names and accession ID are in the supplementary table 2. For the run parameters, multiple sequence alignment (MSA) files were uploaded. MSA was performed using the MUSCLE algorithm embedded in MEGA 11(version 11.0.13) (ref) with default parameters. The MSA file was uploaded in ConSurf and the *S.elongatus* sequence was used as the query for both the FNR and 6-PGD. For the evolutionary conservation calculation in ConSurf the Bayesian method was used and evolutionary substitution model was given as default option.

### Conservation of amino acid-ligand interaction across species

PDB structure of *Nostoc sp. PCC 7119* and *Homo sapiens* were used to determine amino acid ligand interaction of FNR and 6PGD with their ligand Nicotinamide-adenine-dinucleotide phosphate. The missing hydrogen atoms were added to the pdb structure using Protoss (Lippert and Rarey, 2009; Bietz et al., 2014) in ProteinsPlus web service (https://proteins.plus/ ). The protonated files were further used to determine the molecular interactions between protein and ligand from PoseView and poseEdit (Diedrich et al., 2023). MSA was performed as aforementioned and the interacting residues were mapped and checked for conservation across species.

## Data availability

The mass spectrometry proteomics raw data have been deposited to the ProteomeXchange Consortium via the PRIDE(Perez-Riverol et al., 2019) partner repository with the dataset identifier PXD068612. Along with the raw data we have also uploaded the excel files with all details. The transcriptomics raw data have been deposited to NCBI SRA under the BioProject ID: PRJNA1269807 with the dataset identifier SUB15312724.

## Acknowledgements

The authors thank Dr. Shantanu Sengupta of the National Facility for Biochemical and Genomic Resources (**NFBGR**), IGIB for help with SWATH-MS sequencing.

## Funding

This study was supported by the Science and Engineering Research Board, Government of India (SERB grant SRG/2019/000364) and Department of Biotechnology, Government of India (DBT grant BT/PR32511/BRB/10/1803/2019), SR acknowledges the Annual Research Grant from Ashoka University. DBJ was supported by a PhD fellowship from Ashoka University.

## Author contributions

**Anudarsh S.:** Experimental designing; data curation; software; experimentation - performing; formal analysis; validation; investigation; visualization; methodology; writing –review and editing. **Dinesh Balasaheb Jadhav:** Some sample collection, some formal analysis. **Sougata Roy:** Conceiving the original idea; experiment designing; resources; supervision; project administration; formal analysis-some: funding acquisition; writing – original draft; writing – review and editing.

## Disclosure and competing interests’ statement

The authors declare that they have no conflict of interest.

## Notes

### Competing Interest Statement

The authors have declared no competing interest.

