## Supplementary material for "RNA-Binding moonlighting function of metabolic enzymes reveals deep evolutionary roots in Cyanobacteria": Supplentary Figures

S1

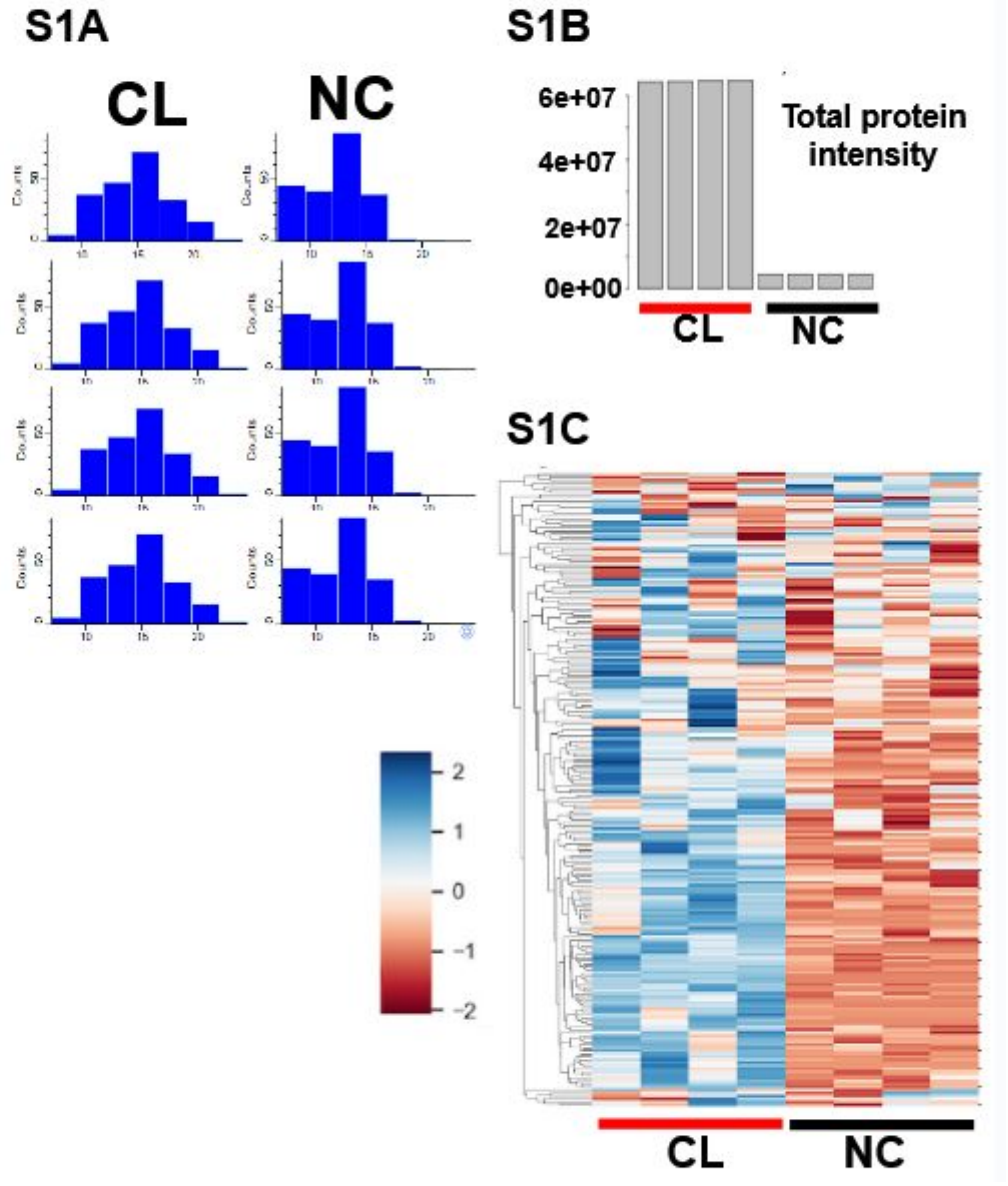


**Supplementary figure S1: CL and NC data quality S1A)** Histogram illustrating the distribution of the protein relative intensity. The left side represents 4 replicates from the CL samples and the right side represents the 4 replicates from the NC samples. **S1B)** Histogram representing the relative intensity of the peptides from CL and the NC samples. **S1C)** Heat map showing the Z normalized relative protein intensity. High protein intensity is represented by blue colour and the low intensity are represented by red.

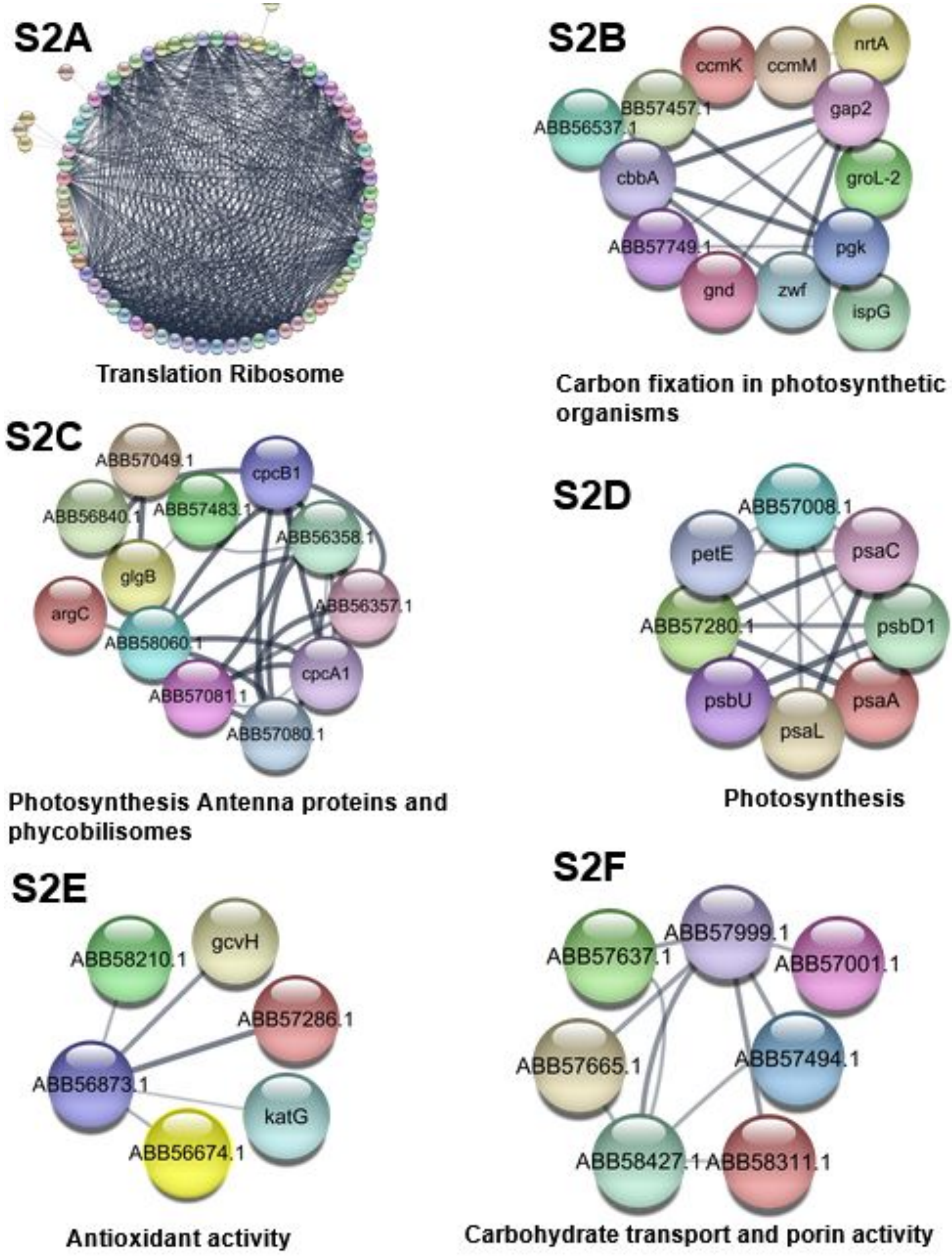


**Supplementary figure S2: RNA-binding protein interaction network.** All the significant 136 cyRBPs were queried on the STRING(https://string-db.org/) using multiple protein search option. The protein interaction network generated was clustered using K-means clustering with node number as eight. Following which all the generated network were transferred to Cytoscape for visualisation. **(S2A)** The translation and ribosome cluster with 73 nodes and have the PPI enrichment p-value <1.0e-16. **(S2B)** The carbon fixation in photosynthetic organism cluster with 13 nodes and have a PPI enrichment p-value <1.0e-16. **(S2C)** Photosynthesis Antenna proteins and phycobilisomes have clustered with 12 nodes and have PPI enrichment p-value< 1.0e-16. **(S2D)** The photosynthesis cluster have eight nodes and have PPI enrichment p-value < 1.0e-16. **(S2E)** The antioxidant activity has clustered with 6 nodes and have PPI enrichment p-value 5.92e-08. **(S2F)** Carbohydrate transport and porin activity have clustered with 7 nodes and have PPI enrichment p-value 2.07e-06.

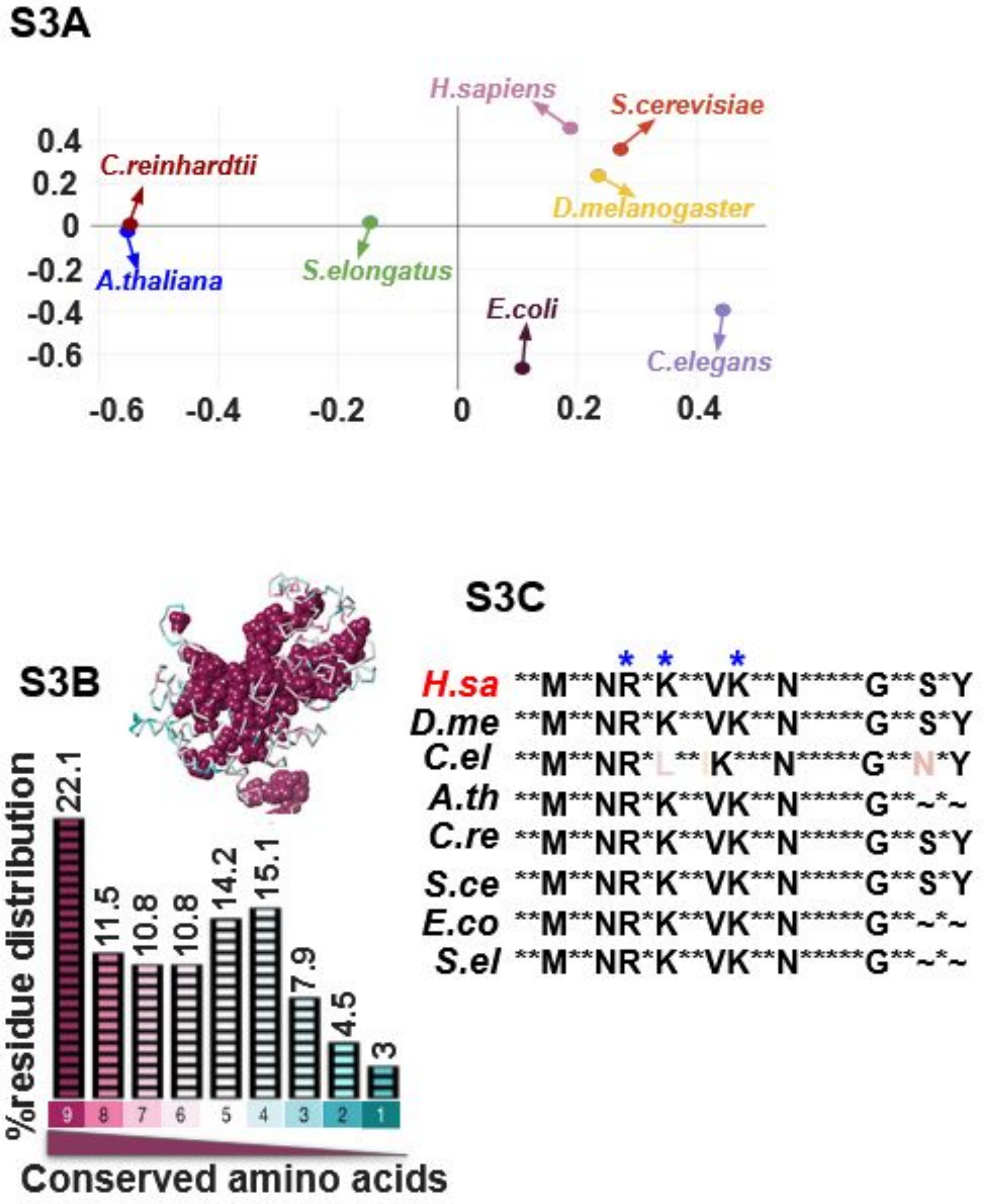


**Supplementary Figure 3: Structural and sequence analysis of the RBP 6-phosphogluconate dehydrogenase, decarboxylating (6PGD) . (S3A)** The correspondence analysis plot shows an all-against-all structural comparison of 6PGD from eight representative species using DALI. Each point corresponds to one protein structure. The relative distances between points reflect structural similarity, calculated from pairwise DALI Z-scores. **(S3B)** The histogram represents the amino acid conservation profile of 6PGD from 28 different different species using ConSurf web server. The conservation score is estimated based on evolutionary conservation of each residue from the uploaded input multiple sequence alignment. The multiple sequence alignment for this was performed using MUSCLE alignment option from MEGA 11(version 11.0.13) with default parameters. Conservation scores were calculated with Bayesian method and amino acid substitution model was chosen by default. **(5 graphical)** The 3D structure of the 6PGD is colour coded based on their conservation scores. Here only the amino acids with conservation scores of 9 are shown as balls and rest of the colour coded sticks represent the following scores of conservations. **(S3C)** The amino acids which interact with the ligand Nicotinamide-adenine-dinucleotide phosphate from the pdb structure (pdb 2jkv) was determined using both PoseView and PoseEdit(https://proteins.plus/).From the interacting ones only the conserved cationic amino acids are highlighted with a blue star. Sequence alignment for this was performed using MEGA11(version 11.0.13) using default parameters.
